# An unexpected specialization of the active zone scaffold RIM at high release synapses

**DOI:** 10.64898/2026.03.20.713218

**Authors:** Rebecca Stark, Prapti Patel, Wanying Dong, Caleb Dehn, Dion Dickman

## Abstract

The active zone scaffold RIM is canonically viewed as an obligate, universal pillar of the neurotransmitter release machinery. However, whether RIM is strictly required across diverse synapse subtypes with distinct release probabilities has remained an important unresolved question. Here, we report a fundamental revision of this model: RIM is not a constitutive necessity for baseline transmission, but rather a specialized “gain factor” selectively deployed to empower high-release synapses. Utilizing botulinum neurotoxin-based silencing to isolate convergent inputs at the Drosophila neuromuscular junction, we demonstrate that while RIM is essential for high-fidelity transmission at phasic (MN-Is) synapses, it is largely dispensable at low-release tonic (MN-Ib) synapses. This input specificity extends into plasticity: RIM is required for acute presynaptic homeostatic potentiation (PHP) at phasic inputs but is dispensable for the chronic maintenance of PHP at tonic inputs. Mechanistically, super-resolution imaging reveals that RIM is positioned significantly closer to CaV2 Ca^2+^ channel nanodomains at phasic synapses. During acute plasticity, RIM coordinates the homeostatic compaction of Ca_V_2 channel clusters and drives the rapid expansion of the readily releasable vesicle pool. Our results revise the canonical view of active zone architecture, demonstrating that rather than serving as a static, uniform structural anchor, RIM functions as a tunable, dynamic module deployed to shape the performance and plasticity of high-demand synaptic connections.

## INTRODUCTION

Neural circuits rely on the precise routing of information through vast networks of synapses that exhibit striking diversity in structural and functional properties. A single postsynaptic neuron may receive thousands of convergent inputs, yet it must independently tune specific connections to distinct “set points” of synaptic strength, plasticity, and neuromodulatory sensitivity (Biederer et al., 2017; Dittman & Ryan, 2019; Nusser, 2018; van Oostrum & Schuman, 2025). This input-specific heterogeneity allows circuits to encode complex information — filtering high-frequency signals at some inputs while acting as high-fidelity relays at others. The presynaptic active zone (AZ) is a primary locus for this specialized transformation. Defined by a conserved nanostructure of protein scaffolds, the AZ controls neurotransmission by organizing voltage-gated Ca^2+^ channels (Ca_V_2), delineating vesicle release sites, and positioning priming factors such as Unc13 to couple Ca^2+^ influx to vesicle fusion (Emperador-Melero & Kaeser, 2020; Haucke et al., 2011). While the molecular “parts list” of the AZ is largely conserved across species, how these components are differentially configured to generate the wide dynamic range of synaptic strength observed across diverse inputs – particularly at connections facing high activity demands – and how they are subsequently modified by plasticity, remains poorly understood.

The multi-domain scaffold RIM (Rab3-interacting molecule) is widely regarded as the central organizer of the active zone. RIM proteins contain multiple conserved interaction domains – including an N-terminal zinc finger flanked by α-helices, a central PDZ domain, and C-terminal C2 domains separated by a proline-rich motif – that form a hub connecting virtually all other AZ components (Kaeser et al., 2011; Südhof, 2012). Extensive biochemical and genetic studies in mammals and invertebrates have established RIM as a key component that anchors the release machinery: RIMs directly interact with Rab3 to recruit synaptic vesicles, bind Unc13/Munc13 via their N-terminal zinc finger to drive vesicle priming, and physically tether Ca_V_2 channels to the AZ via interactions with RIM-Binding Protein (RIM-BP) and a direct PDZ domain-channel C-terminus interaction (Acuna et al., 2016; Brockmann et al., 2020; Deng et al., 2011; Emperador-Melero et al., 2024; Han et al., 2011; Kaeser et al., 2011; Koushika et al., 2001; Südhof, 2012). Importantly, these functions are executed by distinct, modular domains: the zinc finger autonomously activates priming by disrupting autoinhibitory Munc13 homodimerization, while a separate PDZ-RIM-BP module mediates Ca^2+^ channel tethering (Deng et al., 2011; Emperador-Melero et al., 2024; Kaeser et al., 2011). Through this multivalent architecture, RIM simultaneously organizes the three core prerequisites for fast neurotransmitter release: a primed vesicle pool, clustered Ca^2+^ channels, and tight spatial coupling between them, supporting the prevailing view that RIM is an obligate, universal component of all chemical synapses.

Consequently, the loss of RIM generally leads to a reduction in Ca^2+^ channel density and impaired synaptic vesicle fusion (Graf et al., 2012; Kaeser et al., 2011; Schoch et al., 2002). However, the assumption that RIM functions as an obligatory, uniform scaffold at all chemical synapses is being challenged. Emerging evidence suggests that RIM requirements may vary significantly between synapse subtypes — for instance, between excitatory and inhibitory synapses in the hippocampus, or in the differential regulation of dense-core vesicle release at neuromodulatory terminals (Banerjee et al., 2022; Brockmann et al., 2020; Emperador-Melero et al., 2024; C. Liu et al., 2018; H. Liu et al., 2019; Persoon et al., 2019; Robinson et al., 2019).

These findings suggest that rather than serving as a static “brick” in the active zone wall, RIM may function as a tunable variable, differentially deployed to match the specific performance requirements of a synapse. Yet, a rigorous mechanistic dissection of this specialization has been hindered by the difficulty of assessing RIM function at distinct, convergent inputs within an intact circuit and by the genetic redundancy of *rim* genes in mammals.

The Drosophila neuromuscular junction (NMJ) offers a powerful genetic system to resolve these questions about input-specific RIM functions. At the fly NMJ, a single *rim* gene encodes the conserved functions of stabilizing Ca_V_2 channels and recruiting the readily releasable pool (RRP) (Graf et al., 2012). Furthermore, subsequent work identified RIM as a critical effector of presynaptic homeostatic potentiation (PHP), a form of adaptive plasticity that upregulates presynaptic neurotransmitter release following the genetic loss or pharmacological blockade of postsynaptic receptors (Müller et al., 2012). Specifically, RIM was reported to mediate the homeostatic expansion of the RRP required to restore synaptic strength (Müller et al., 2012). However, a critical confound exists in these foundational studies: electrophysiological recordings were performed on muscles co-innervated by two distinct motor neurons — the tonic “Type Ib” (MN-Ib) and the phasic “Type Is” (MN-Is) — which differ profoundly in their basal release properties, short-term plasticity, and AZ nanoarchitecture (Aponte-Santiago & Littleton, 2020; Chien et al., 2025; He et al., 2023; He & Dickman, 2025; Jetti et al., 2023; Medeiros et al., 2023). In addition, most imaging characterizing *rim* mutants was performed primarily at MN-Ib terminals, leaving potential differences at MN-Is unexamined and thereby introducing a disconnect between blended electrophysiological phenotypes and imaging analyses focused predominantly on MN-Ib. Because standard electrophysiological recordings represent an ambiguous composite of these dramatically divergent inputs, and prior studies did not separate them, it has remained unknown whether RIM is indeed universally required for basal transmission and plasticity at both inputs, or whether the phenotypes observed in *rim* mutants reflect the selective failure of one synaptic subtype.

Here, we use botulinum neurotoxin–based genetic silencing to isolate transmission from tonic MN-Ib and phasic MN-Is inputs and rigorously test the synapse-type specificity of RIM. We uncover a striking dichotomy: while RIM is essential for high-fidelity release at phasic MN-Is synapses, basal transmission at tonic MN-Ib is largely unchanged in *rim* null mutants. This input specificity extends into plasticity, where RIM is required for the acute expression of PHP at MN-Is but is dispensable for the chronic maintenance of PHP at MN-Ib. Mechanistically, endogenous tagging and super-resolution imaging reveal that RIM is differentially organized at the nanoscale, positioned closer to Ca_V_2 clusters at MN-Is where it maintains tight Ca_V_2–vesicle coupling and supports rapid RRP expansion. Furthermore, during acute plasticity, RIM coordinates the homeostatic compaction of Ca_V_2 channel clusters to recruit additional vesicles for fusion. These findings revise the canonical view of RIM, revealing it not as a constitutive necessity for all release, but as a specialized, tunable gain module deployed to boost synaptic strength and plasticity specifically at high-demand connections.

## RESULTS

### RIM selectively promotes neurotransmitter release at phasic MN-Is synapses

To determine whether the AZ scaffold RIM contributes uniformly to synaptic transmission, we exploited the Drosophila larval NMJ, where the tonic MN-Ib and phasic MN-Is coinnervate muscle 6, exhibiting dramatically distinct release properties (**Fig. 1A,B**). Specifically, tonic MN-Ib terminals exhibit lower release probability and facilitate upon repeated stimulation, while phasic MN-Is terminals possess a higher P_r_ and depress during repeated stimulation (He et al., 2023; Lu et al., 2016; Newman et al., 2017). We first generated new targeted *rim* null mutant alleles in an isozygous genetic background (*w^1118^*), ensuring consistency across all genotypes evaluated in this study. Using CRISPR/Cas9 gene editing, we isolated two new alleles (*rim^RS1^* and *rim^RS2^*) (**Fig. S2A**), confirming the expected anatomic (Table S1) and electrophysiologic phenotypes reported previously (Graf et al., 2012; Müller et al., 2012). These phenotypes included normal miniature events and a ∼50% reduction in evoked responses during blended, co-stimulated (Is+Ib) transmission at NMJs (**Fig. 1C-E**). Given that both *rim* alleles exhibited similar expected phenotypes, we focused exclusively on *rim^RS1^* for the remainder of our experiments.

**Figure 1:**
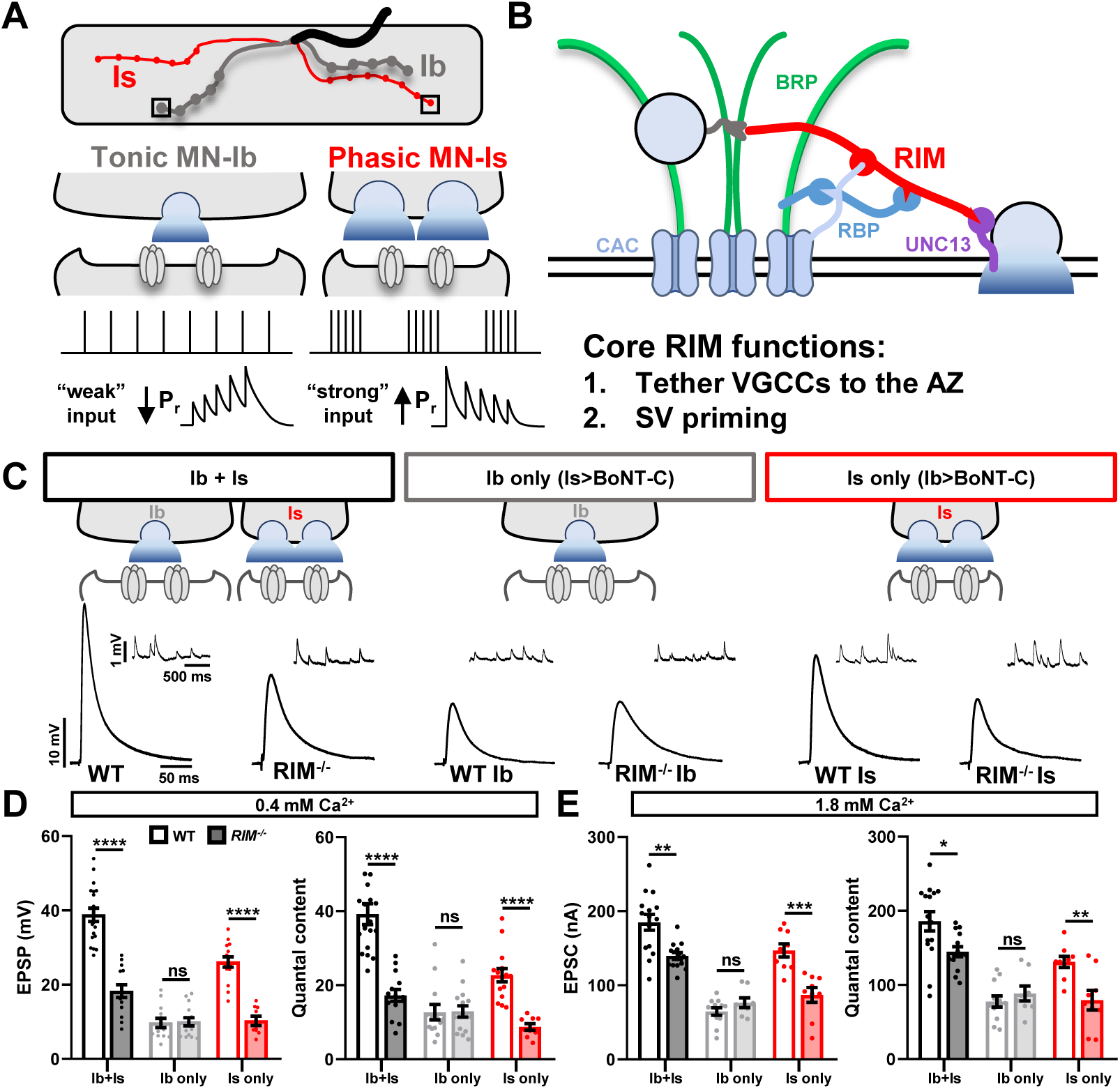
RIM selectively promotes neurotransmitter release at phasic MN-Is synapses. **(A)** Schematic of the Drosophila NMJ, where motor neurons Ib and Is co-innervate abdominal muscle 6. Tonic MN-Ib synapses function with low release properties, while phasic MN-Is operate with high release characteristics. **(B)** Schematic of the active zone, highlighting key interactions of RIM with associated release machinery. **(C)** Representative mEPSP and EPSP traces at baseline (wild type; *w^1118^*) and in *rim* mutants (*w;;rim^RS1^*) during co-stimulation (Ib+Is) or following input-specific silencing with BoNT-C (MN-Ib or MN-Is only). Evoked amplitude is unchanged at MN-Ib in the absence of *rim*, while transmission from MN-Is is reduced by ∼60%. **(D,E)** Quantification of evoked responses and quantal content values at 0.4 mM (D) and 1.8 mM (E) extracellular [Ca^2+^], revealing selective deficits exclusively at *rim* MN-Is. Error bars indicate ±SEM. Additional statistical details are provided in Table S1.

To resolve the input-specific requirements of *rim* at tonic versus phasic synapses, we isolated MN-Ib or MN-Is transmission via the targeted expression of botulinum neurotoxin C (BoNT-C), which silences transmission at the convergent input (i.e., MN-Is>BoNT-C to isolate MN-Ib, referred to as “Ib only”, and vice versa for “Is only”) (Han et al., 2022; He et al., 2023).

Unexpectedly, at isolated MN-Ib synapses in *rim* mutants, we observed no significant change in EPSP responses in *rim* mutants compared to wild type controls at reduced extracellular Ca^2+^ (0.4 mM) (**Fig. 1C,D**), nor did EPSC amplitudes differ under higher Ca^2+^ conditions (**Fig. 1E**; **S1A,B**). We probed these release properties in further detail at isolated MN-Ib synapses, finding no significant differences in EPSC amplitude or quantal content across a wide range of extracellular Ca^2+^ conditions (0.4 – 6 mM). Furthermore, failure analysis and paired-pulse facilitation revealed no differences in release probability for *rim* mutants at MN-Ib (**Fig. S1A-H**). These surprising findings implied that all observed defects in neurotransmission following the loss of *rim* derive exclusively from alterations in phasic MN-Is release properties.

Indeed, at isolated MN-Is synapses, the loss of *rim* reduced release by a sufficient magnitude to explain the defects previously reported, which were formerly interpreted as resulting from combined MN-Ib and MN-Is decrements. Specifically, both EPSP and EPSC amplitudes were reduced across all extracellular Ca^2+^ concentrations evaluated in *rim* MN-Is synapses (**Fig. 1C-E**; **S1C,D**), while failure and paired-pulse analyses confirmed a significantly decreased P_r_ (**Fig. S1E-H**). Together, these data demonstrate that *rim* is selectively required to support high release probability transmission at phasic MN-Is synapses, whereas tonic MN-Ib transmission remains essentially unperturbed in its absence.

### RIM is enriched at MN-Is active zones

We next asked whether differences in RIM abundance or nanoscale organization might explain its apparent importance in promoting neurotransmitter release selectively at phasic MN-Is synapses. To address this, we generated an endogenously tagged *rim* allele using a short ALFA tag (*rim^ALFA^*), targeting a region near the N-terminal Zinc Finger domain previously validated in a recent study (Mrestani et al., 2023) (**Fig. S2A**). Characterization of *rim^ALFA^* larvae confirmed normal synaptic growth, transmission, and PHP plasticity expression compared to controls (**Fig. S2B–F**), validating that RIM was unperturbed by the tag and justifying its use for localization analyses.

We first systematically examined six central AZ components across MN-Ib and -Is terminals to determine whether RIM or any other proteins exhibited differential localization or abundance. Confocal imaging of RIM^ALFA^ alongside BRP, RBP, CAC (*cac^sfGFP^*), Unc13A, and Unc13B revealed no significant differences in CAC, Unc13A, or Unc13B levels across the two input types (**Fig. 2A,C**). Additionally, we confirmed previously reported reductions in intensity for BRP and RBP at Is synapses relative to Ib (**Fig. 2A,B**) (He et al., 2023; Medeiros et al., 2023). However, RIM intensity was significantly higher at MN-Is AZ compared to MN-Ib (**Fig. 2A,B**), which is notable as it represents the only AZ component examined that enriched at Is over Ib synapses. Next, we utilized super-resolution (STED) microscopy to resolve the nanoscale organization of these components. Within a single AZ labeled by the “T-bar” scaffold BRP, we observed approximately two RIM puncta at both MN-Ib and MN-Is, indicating no significant difference in cluster number (**Fig. 2D,E**). However, when we imaged RIM and Ca_V_2 (CAC) channels, we observed a significantly tighter distances between RIM and CAC at MN-Is compared to MN-Ib (**Fig. 2F,G**). Thus, RIM is not only enriched but also positioned closer to Ca_V_2 nanodomains at phasic MN-Is synapses.

**Figure 2:**
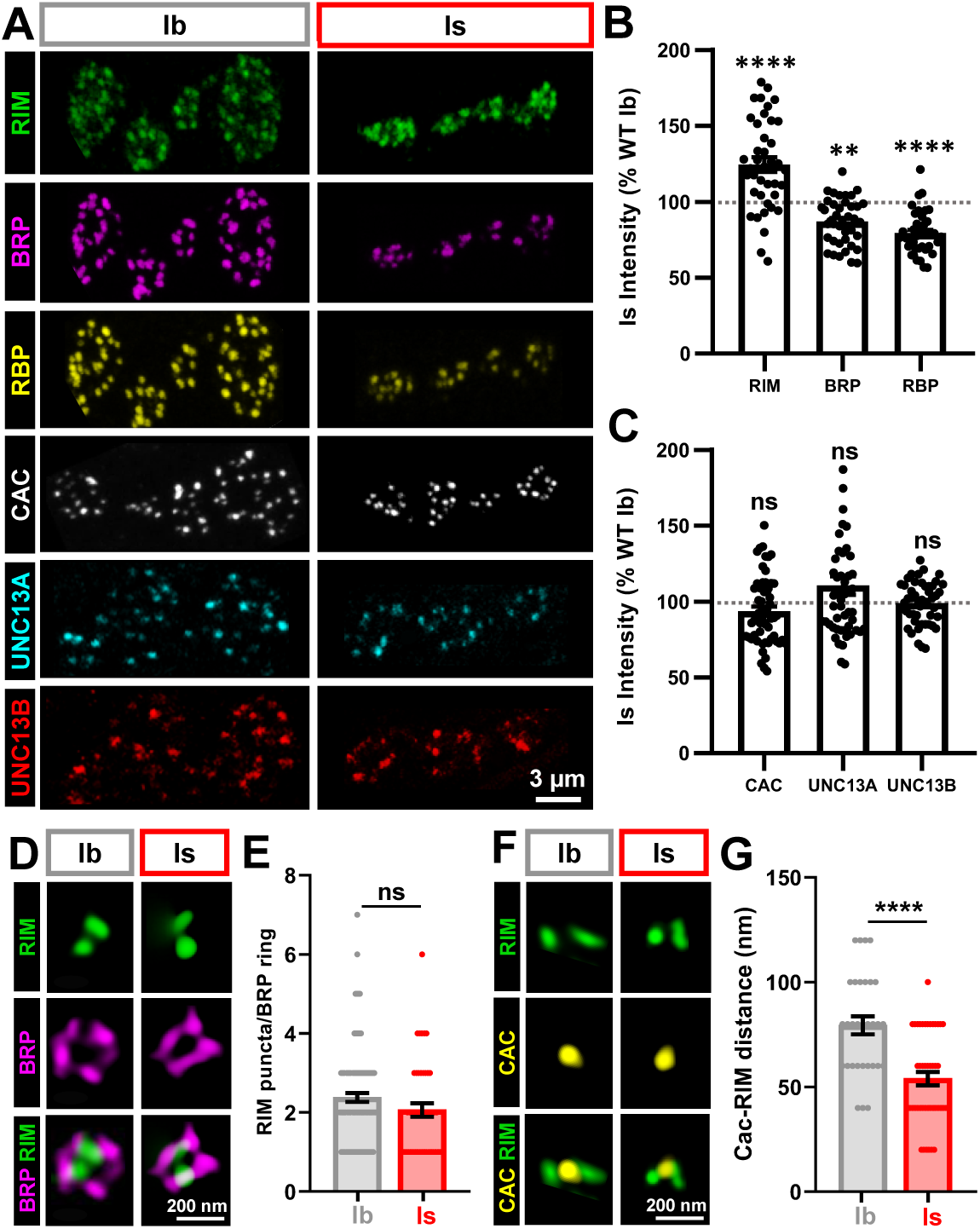
RIM is uniquely abundant at MN-Is active zones, localizing close to Ca_V_2 channels. **(A)** Representative confocal images of muscle 6 NMJs immunostained for the AZ components RIM (ALFA; *w;;rim^ALFA^*), BRP, RBP, CAC (GFP; *Cac^sfGFP^*^-N^) at MN-Ib and -Is. **(B)** Quantification of mean immunofluorescence intensity for the indicated antigens at MN-Is, normalized as a percentage of MN-Ib. RIM intensity is uniquely higher at MN-Is, whereas BRP and RBP levels are higher at MN-Ib. **(C)** Quantification as in (B) for CAC, Unc13A, and Unc13B, indicating similar intensities between MN-Ib and -Is. **(D)** Representative STED images of single AZs immunostained for BRP and RIM (ALFA) at MN-Ib and MN-Is. **(E)** Quantification of RIM puncta per BRP ring. **(F)** Representative STED images of single AZs immunostained for CAC (GFP) and RIM (ALFA). **(G)** Quantification of CAC-RIM distance, indicating closer nano-organization at MN-Is compared to MN-Ib. Error bars indicate ±SEM. Additional statistical details are shown in Table S1.

We next sought to determine how active zone (AZ) organization at MN-Ib and MN-Is is impacted by loss of *rim*, given that RIM is a conserved AZ scaffold implicated in vesicle priming and Ca²⁺ channel organization at synapses (Graf et al., 2012; Han et al., 2011; Kaeser et al., 2011). We systematically assessed the remaining five AZ components (BRP, RBP, CAC, Unc13A, and Unc13B) at each input using confocal microscopy. Consistent with previous studies showing that RIM contributes to Ca²⁺ channel recruitment and release-site organization at the Drosophila NMJ (Graf et al., 2012; Müller et al., 2012), we observed a small but expected reduction in CAC levels, while BRP and RBP levels were unchanged at both inputs (**Fig. 3A,B,E,F).**

**Figure 3:**
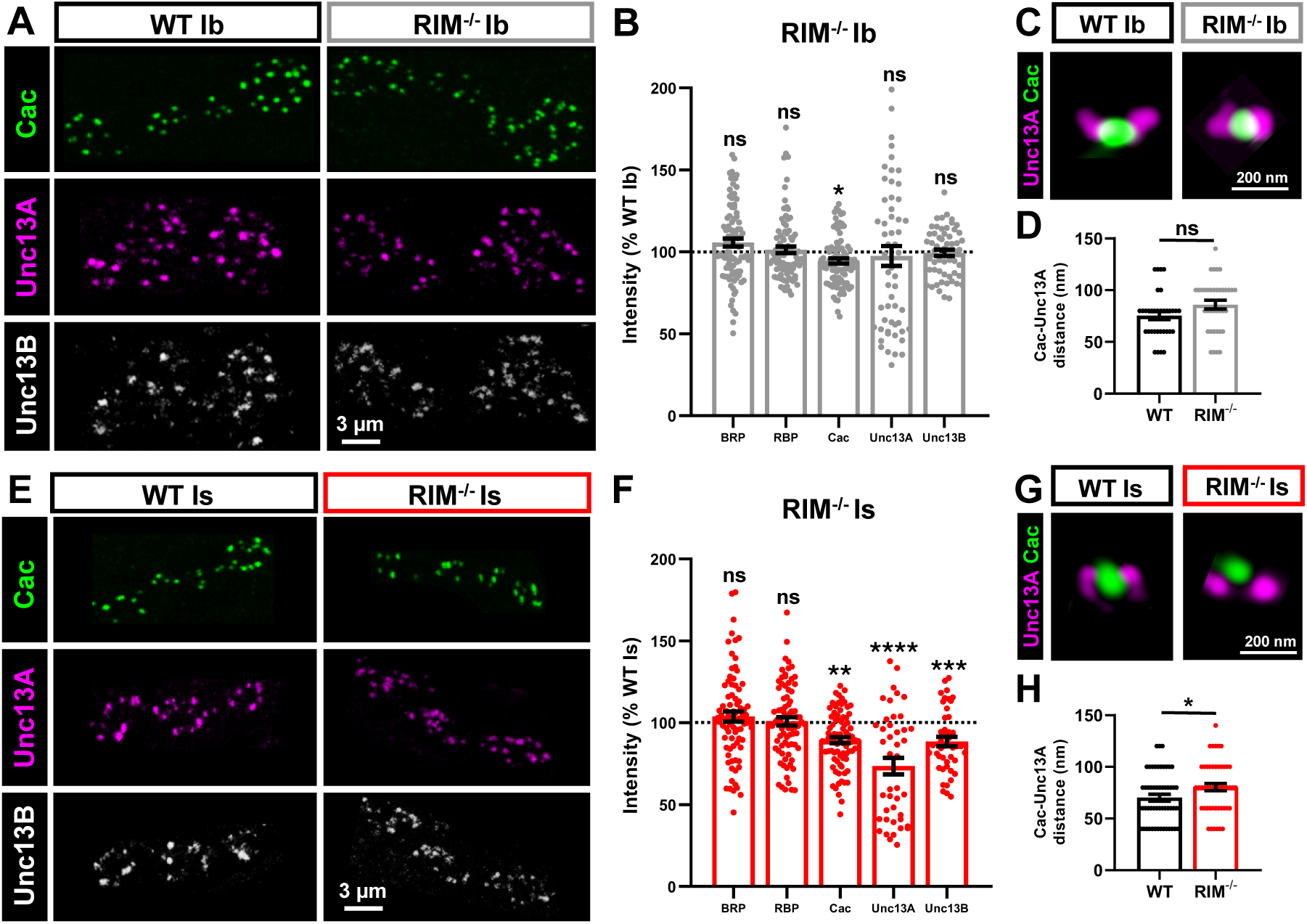
RIM is required for the accumulation of CAC and positioning of Unc13A near Ca_V_2 channels. **(A)** Representative confocal images of MN-Ib NMJs immunostained for CAC, Unc13A, and Unc13B in wild type and *rim* mutants. **(B)** Quantification of mean immunofluorescence intensity for the indicated antigens in *rim* mutants as a percentage of wild type. Loss of *rim* leads to a small, selective decrease in CAC levels. **(C)** Representative STED images of single AZs immunostained for Unc13A and CAC at MN-Ib in wild-type and *rim* mutants. **(D)** Quantification of Cac–Unc13A distances at MN-Ib NMJs, with no significant change observed in the absence of *rim*. **(E-H)** Corresponding data for MN-Is. At MN-Is, loss of *rim* significantly reduces CAC, Unc13A, and Unc13B levels, and causes a small but significant increase in Cac-Unc13A distance. Error bars indicate ±SEM. Additional statistical details are provided in Table S1.

We next examined the vesicle-priming factors Unc13A and Unc13B, which are differentially positioned at release sites and contribute to synaptic vesicle–Ca²⁺ channel coupling at the NMJ (Böhme et al., 2016). While no changes were detected for either Unc13 isoform at MN-Ib synapses, loss of *rim* resulted in a moderate reduction in Unc13B and a substantial diminution of Unc13A specifically at MN-Is terminals (**Fig. 3A,B,E,F**).

Given the pronounced reduction of Unc13A selectively at MN-Is AZs, we next asked whether the spatial relationship between Unc13A and Ca²⁺ channels was altered in the absence of RIM. Using STED imaging, which has previously revealed the nanoscale organization of Unc13 isoforms relative to Ca²⁺ channels at Drosophila active zones (Böhme et al., 2016), we measured CAC–Unc13A distances in *rim* mutants. While no significant difference was observed at MN-Ib AZs, Unc13A was positioned significantly further from CAC at MN-Is synapses in the absence of *rim* (**Fig. 3C,D,G,H**). Together, these results indicate that RIM is selectively required at phasic MN-Is synapses to stabilize Unc13A in close proximity to Ca²⁺ channels and maintain tight vesicle–channel coupling.

### RIM selectively promotes presynaptic homeostatic potentiation

We next focused on clarifying whether the unanticipated, input-specific functions of *rim* at phasic MN-Is synapses extend to roles in plasticity as well. Presynaptic homeostatic potentiation (PHP) is a well-studied form of plasticity modeled at the Drosophila NMJ, where pharmacological blockade or genetic loss of postsynaptic glutamate receptors triggers a retrograde, adaptive enhancement in presynaptic release to maintain stable synaptic strength (Davis & Müller, 2015; Frank et al., 2006, 2020; Goel & Dickman, 2021). A previous study identified *rim* as essential for acute PHP (Müller et al., 2012), which was interpreted as resulting from a genetically separable role in Ca^2+^ influx and the expansion of the readily releasable vesicle pool (RRP). Because the blended physiological responses from MN-Ib and -Is confounded this interpretation, an alternative hypothesis is that the loss of *rim* did not actually illuminate genetically separable processes in PHP, but rather that these processes are dictated by input-specific functions during PHP. Indeed, a recent study demonstrated that two forms of PHP utilize distinct mechanisms, and target distinct inputs, to achieve stable NMJ strength (Chien et al., 2025). “Chronic PHP”, induced through the genetic loss of the *GluRIIA* subunit, selectively drives enhanced presynaptic Ca^2+^ influx at MN-Ib to promote enhanced vesicle release and maintain stable synaptic strength (schematized in **Fig. 4A**). In contrast, “acute PHP”, induced through the pharmacological application of the glutamate receptor antagonist philanthotoxin (PhTx), selectively targets MN-Is for homeostatic modulation, mobilizing an expanded RRP pools to drive the increased transmitter release that stabilizes synaptic strength (**Fig. 4E**). Given these critical new insights into the input-specific mechanisms orchestrating PHP, we focused on accurately assessing *rim’s* functions across these distinct forms of plasticity.

**Figure 4:**
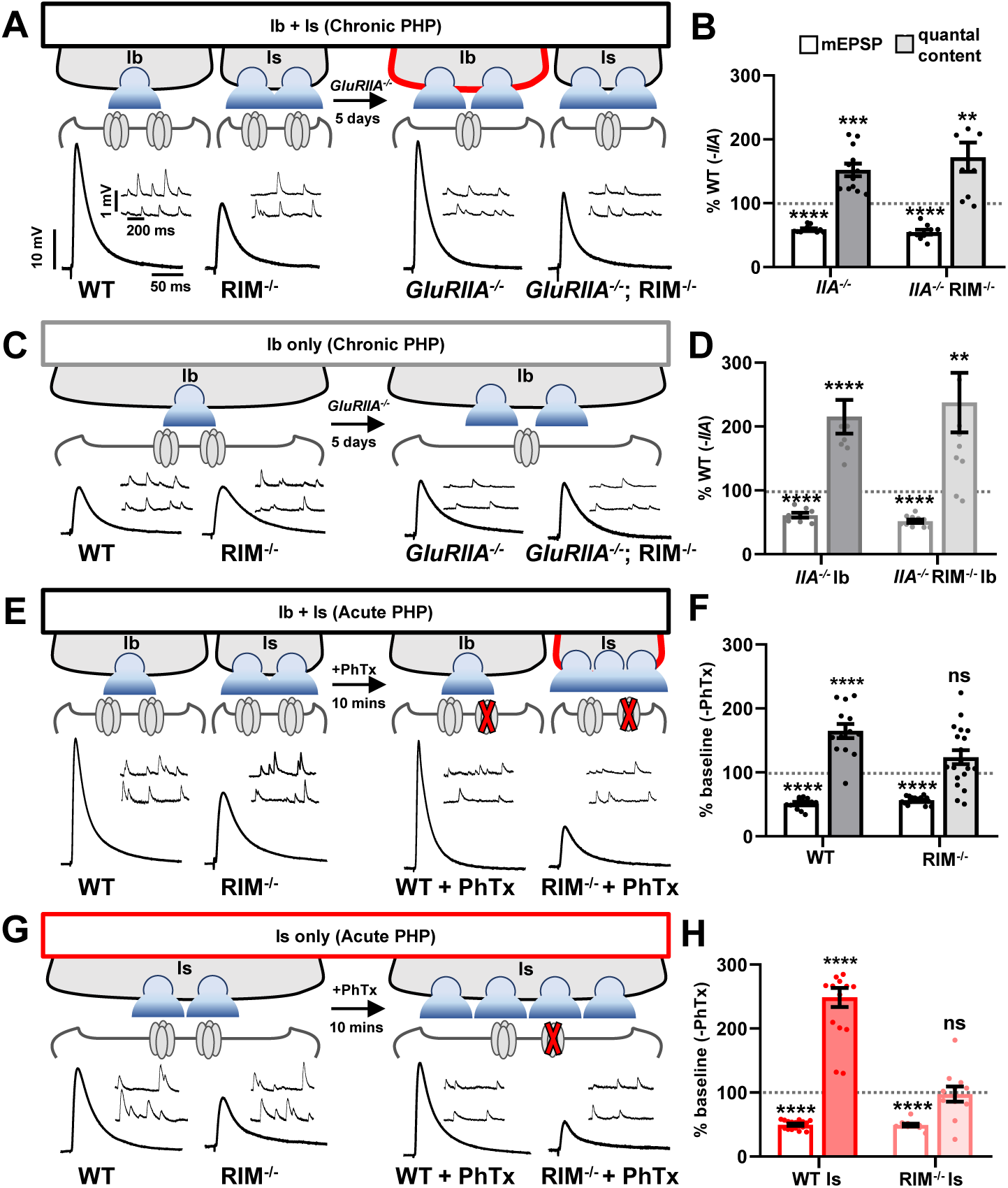
RIM selectively promotes acute PHP at MN-Is. **(A)** Representative EPSP traces for baseline (WT: *w^1118^*), chronic PHP (GluRIIA^−/−^: *w*;*GluRIIA^PV3^*), *rim^-/-^* (w;;*rim^-/-^*), and chronic PHP in *rim ^-/-^* (*w;GluRIIA^PV3^;rim ^-/-^*) NMJs at 0.4 mM extracellular [Ca^2+^]. **(B)** Quantification of mEPSP amplitude and quantal content normalized as a percentage of baseline. Quantal content is significantly enhanced in both WT and *rim^-/-^*, indicating robust chronic PHP expression. **(C)** Representative EPSP traces for isolated MN-Ib following selective expression of BoNT-C in MN-Is. **(D)** Quantification of the data shown in (C), confirming chronic PHP is expressed at MN-Ib lacking *rim*. **(E-F)** Corresponding data for acute PHP (PhTx application) at co-stimulated inputs (Ib+Is). Quantal content does not change in *rim*+PhTx, indicating a block in acute PHP expression. **(G,H)** Corresponding data for acute PHP at isolated MN-Is. Acute PHP is not expressed in isolated *rim* MN-Is NMJs. Error bars indicate ±SEM. Additional statistical details are provided in Table S1.

Whether *rim* is necessary for chronic PHP has previously remained unknown. We therefore generated *GluRIIA*; *rim* double mutants to assess chronic PHP using blended (Ib+Is) physiology. We found that while presynaptic release (quantal content) was robustly enhanced in *GluRIIA* mutants alone, a similar enhancement was observed in *GluRIIA*; *rim* double mutants, demonstrating that *rim* is not required for chronic PHP expression (**Fig. 4A,B**). As expected, in isolated MN-Ib recordings using BoNT-C expression at MN-Is, chronic PHP was robustly expressed despite the loss of *rim* (**Fig. 4C,D**). In contrast, we confirmed that *rim* is required for acute PHP expression when evaluating co-stimulated inputs (**Fig. 4E,F**), as previously reported (Müller et al., 2012). Crucially, PhTx application to isolated *rim* mutant MN-Is terminals revealed that *rim* is strictly necessary at phasic MN-Is for acute PHP expression (**Fig. 4G,H**). Together, these data demonstrate that *rim’s* previously assigned functions in Ca^2+^ channel stabilization and RRP modulation do not genetically separate PHP mechanisms but rather constitute input-specific functions at high P_r_ phasic MN-Is synapses that extend to homeostatic plasticity, proving that *rim* is selectively required for acute PHP at these phasic inputs.

Because acute PHP selectively targets phasic MN-Is synapses by enhancing the RRP (Chien et al., 2025), we next determined whether *rim* was necessary for this specific process. First, we assessed whether *rim* regulates the baseline RRP size at isolated MN-Ib versus Is synapses. To estimate the RRP, we stimulated NMJs at high frequency (60 Hz) under elevated P_r_ conditions (3 mM extracellular Ca^2+^) and plotted the cumulative EPSC. A linear fit was applied from stimulus 19 to 30 and back-extrapolated to time 0 (y-intercept) to estimate the cumulative EPSC (**Fig. 5A,B**). To estimate the RRP from these data, the cumulative EPSC is divided by the respective cell’s quantal size (Goel, Li, et al., 2019). This analysis revealed that the baseline RRP was reduced by approximately 50% at MN-Is in *rim* mutants (**Fig. 5A-C**) but remained unaffected at MN-Ib (Table S1). Following acute PHP induction by PhTx application, control MN-Is exhibited a robust enhancement of RRP size, while in *rim* mutants, the RRP failed to increase (**Fig. 5A-C**), matching previously published results from combined MNs Ib and Is (Müller et al., 2012). Together, these results highlight that *rim* is necessary for both establishing the baseline RRP size and driving its homeostatic expansion specifically at high release phasic MN-Is NMJs.

**Figure 5:**
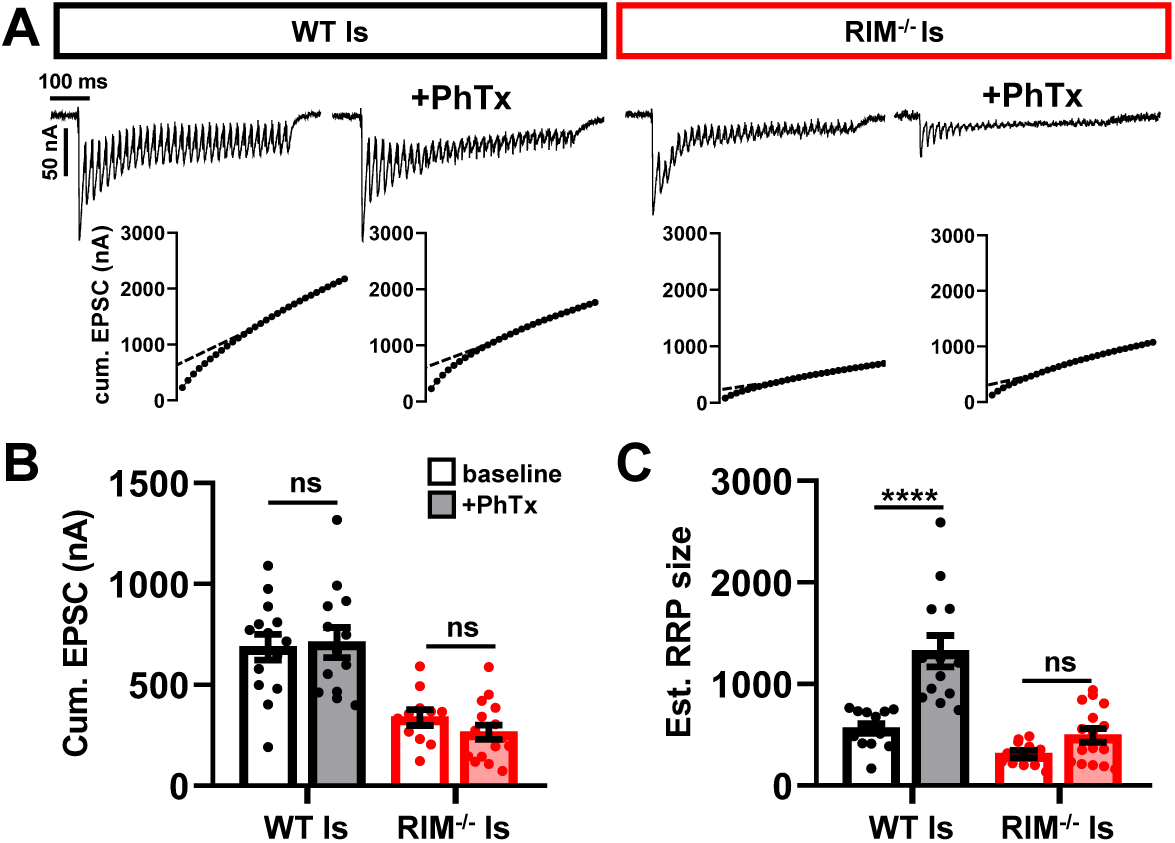
RIM is required for the homeostatic increase in the RRP that enables acute PHP. **(A)** Representative traces of 30 EPSCs from wild-type and *rim* mutants at isolated MN-Is NMJs stimulated at 60 Hz in 3 mM extracellular [Ca^2+^] at baseline and after PhTx application. Bottom: Averaged cumulative EPSC amplitudes. A linear fit applied to the 19^th^-30^th^ stimuli was back extrapolated to time 0. **(B,C)** Quantification of cumulative EPSC (B) and estimated RRP size (C). The characteristic increase in RRP size during acute PHP fails in *rim* mutants. Error bars indicate ±SEM. Additional statistical details are provided in Table S1.

### RIM coordinates the homeostatic compaction of AZ nanostructure

In our final set of experiments, we focused on RIM and the homeostatic re-organization of AZ nanostructure. A variety of previous studies have shown that AZ components at MN-Ib and -Is remodel after both chronic and acute PHP (Böhme et al., 2019; Goel, Dufour Bergeron, et al., 2019; Goel et al., 2017; Gratz et al., 2019; Weyhersmüller et al., 2011). At MN-Is, the AZ proteins BRP, RBP, and CAC increase in intensity following PhTx application (Chien et al., 2025; Medeiros et al., 2023), and super resolution imaging has revealed a “compaction” of AZ nanostructure (Chien et al., 2025; Ghelani et al., 2023; Mrestani et al., 2023), including a compression of CAC channels into a smaller area at AZs (Chien et al., 2025; Ghelani et al., 2023). First, we examined how RIM responds to acute PHP signaling, observing a similar increase in fluorescence intensity at MN-Is AZs following PhTx application (**Fig. 6A,B**). Next, using STED microscopy, we interrogated the homeostatic remodeling of RIM nanostructure. This analysis revealed that while there was no significant change in the number of RIM puncta at MN-Is AZs following acute PHP (**Fig. 6C,D**), the distance between RIM and CAC substantially decreased from ∼60 nm to ∼40 nm (**Fig. 6E,F**). We then confirmed that acute PHP compacts CAC nanostructure at MN-Is AZs, but in the absence of *rim*, CAC channel area does not change (**Fig. 6G,H**), demonstrating that the homeostatic compaction of CAC is gated through a RIM-dependent mechanism. Thus, acute PHP compacts CAC channels at phasic AZs through a RIM-dependent mechanism, where RIM and CAC are re-organized into close proximity.

**Figure 6:**
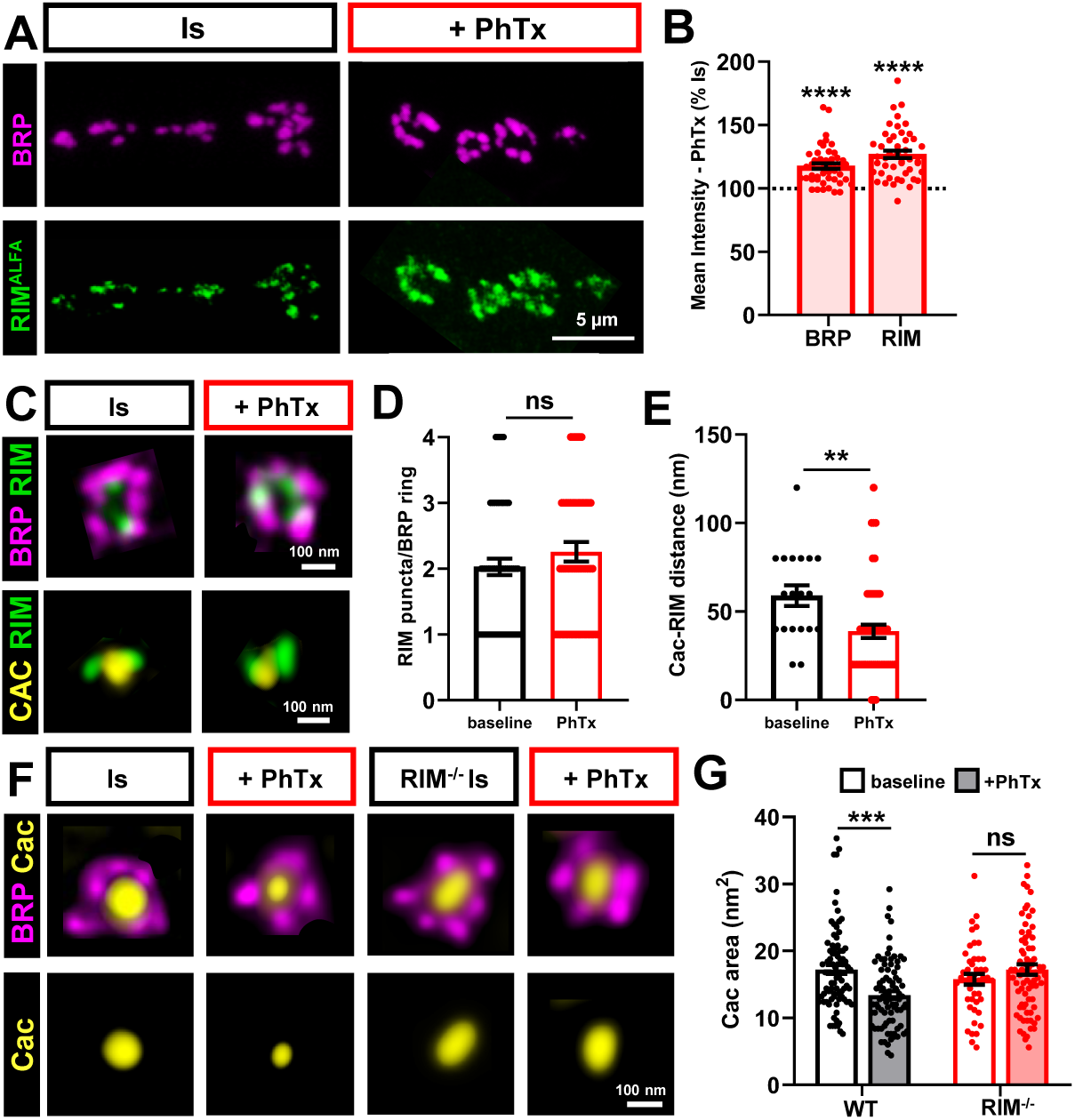
RIM is positioned closer to Ca_V_2 channels and is required for their compaction in acute PHP. **(A)** Representative confocal images of MN-Is NMJs immunostained for BRP and RIM (ALFA) at baseline and after PhTx application. **(B)** Quantification of BRP and RIM intensities after PhTx application as a percentage of baseline, indicating enhanced intensity of both during acute PHP. **(C)** Representative STED images of single AZs immunostained for RIM and BRP or CAC (GFP) in the indicated conditions. **(D)** Quantification of RIM puncta per BRP ring, indicating that the number of RIM puncta does not change during acute PHP. **(E)** Quantification of distance between CAC and RIM puncta at baseline and after PhTx application, showing tighter coupling during acute PHP. **(F)** Representative STED images of single AZs immunostained for BRP and CAC at MN-Is at baseline and after PhTx application in wild type and *rim* mutants. **(G)** Quantification of CAC area after PhTx application as a percentage of baseline. CAC channels compact during acute PHP in wild-type synapses, but fail to compact in *rim* mutants. Error bars indicate ±SEM. Additional statistical details are provided in Table S1.

What adaptive functions would the RIM-dependent compaction of CAC channels serve to promote enhanced neurotransmitter release following acute PHP signaling? One attractive possibility is that the smaller area of CAC channels and associated machinery enables more synaptic vesicles to be located near Ca^2+^ sources to be released during evoked activity. To test this hypothesis, we subjected isolated wild-type and *rim* mutant MN-Is synapses to EGTA-AM incubation at baseline and following PhTx application (**Fig. 7A**). EGTA-AM competes for access to Ca^2+^ diffusing from Ca_V_2 channels with “loosely coupled” synaptic vesicles located a significant distance from the Ca^2+^ source (Kaeser & Regehr, 2014; Meinrenken et al., 2002). At baseline, EGTA-AM application has a marked impact on transmission from *rim* MN-Is synapses compared to wild-type controls, reducing EPSC amplitude more severely (**Fig. 7A,B**); this increase in the proportion of loosely coupled vesicles is consistent with RIM helping to position Unc13A near CAC channels and drive tightly coupled vesicle fusion (Fig. 3G,H). However, EGTA-AM application to PhTx-treated MN-Is synapses led to an additional reduction in EPSC amplitudes in controls, but no change in *rim* mutants (**Fig. 7A,B**). This indicates that acute PHP does indeed recruit additional loosely coupled vesicles for release, and that this occurs through a RIM-dependent mechanism. STED imaging revealed that acute PHP reduces the distance between CAC and Unc13A, and that this enhanced proximity is RIM-dependent (**Fig. 7C,D**). Taken together, these results support a model in which RIM homeostatically re-organizes AZ nanostructure in acute PHP, coordinating the compression of CAC channels and positioning Unc13A – which defines sites for synaptic vesicle fusion – near Ca^2+^ sources to recruit additional loosely coupled vesicles. We present a schematic summarizing these roles for RIM in baseline and homeostatic plasticity specifically at phasic, high release probability synapses (**Fig. 7E**).

**Figure 7:**
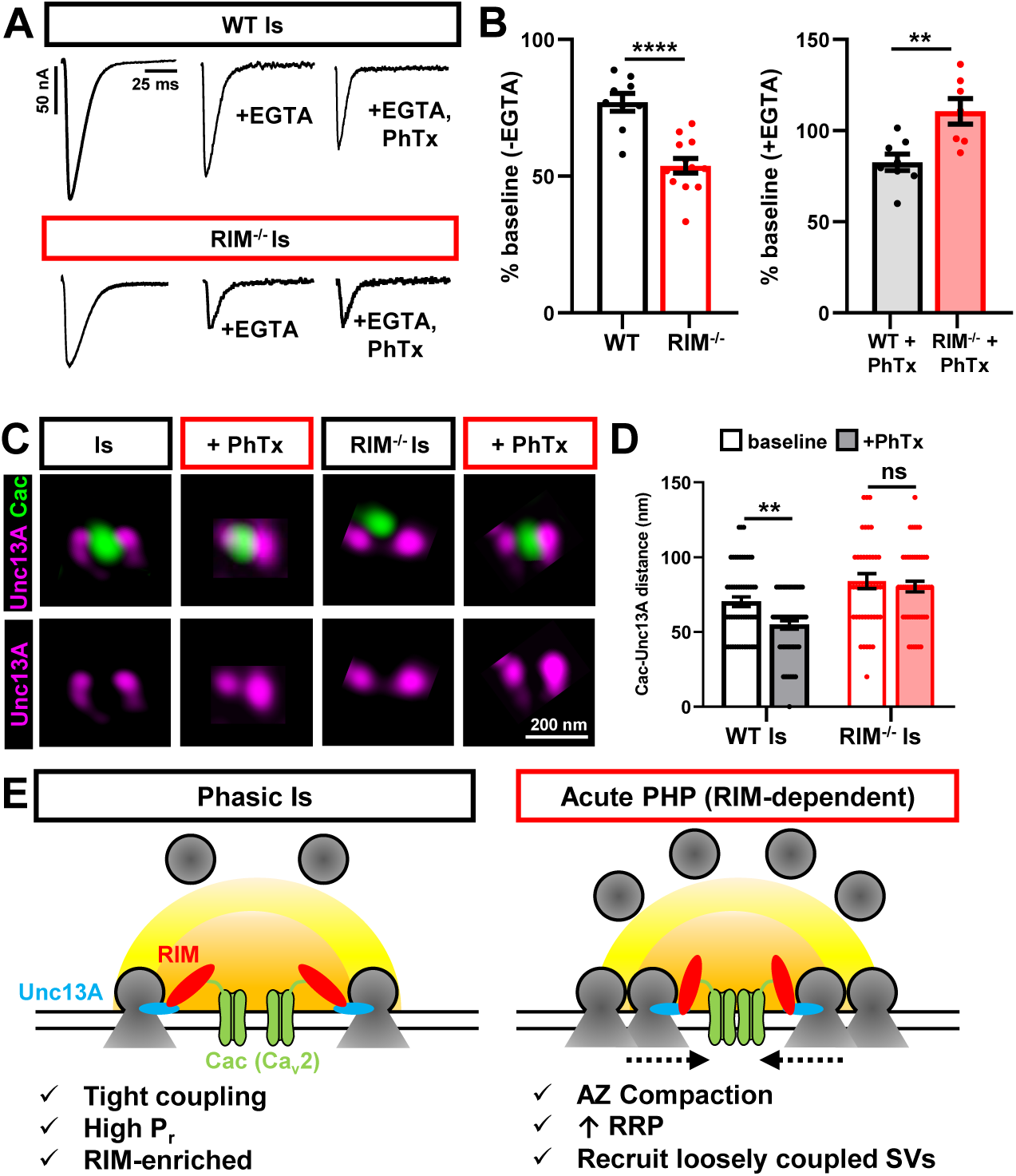
RIM is required for the enhanced Ca^2+^ coupling in acute PHP. **(A)** Representative EPSC traces at 1.8mM [Ca^2+^] from isolated wild-type or *rim* MN-Is synapses at baseline, following EGTA-AM application, and following EGTA-AM with PhTx. **(B)** Quantification of EPSC amplitude after EGTA treatment normalized to baseline (left) or after PhTx application (right). A higher proportion of vesicles are loosely coupled in *rim* mutants at baseline. Following PhTx application, loosely coupled vesicles homeostatically increase in wild-type synapses, but remain unchanged in *rim* mutants. **(C)** Representative STED images of single MN-Is AZs immunostained for CAC and Unc13A at baseline and after PhTx application. **(D)** Quantification of CAC-Unc13A distance, showing that acute PHP drives a RIM-dependent tightening of CAC-Unc13A nano-organization. **(E)** Schematic illustrating the role of RIM in promoting neurotransmission selectively at phasic MN-Is synapses at baseline and during acute PHP plasticity. Error bars indicate ±SEM. Additional statistical details are provided in Table S1.

## DISCUSSION

The AZ scaffold RIM has long been viewed as an obligate, universal pillar of the presynaptic release machinery. However, our input-specific dissection at the Drosophila NMJ reveals a fundamentally different paradigm. By utilizing neurotoxin-based silencing to isolate transmission at convergent inputs, we demonstrate that RIM is not a constitutive necessity for baseline neurotransmitter release, but rather a specialized “gain factor” deployed selectively to empower high-release probability synapses. Specifically, tonic, low-P_r_ synapses function remarkably well without RIM, whereas high-fidelity phasic transmission depends critically on its presence. This necessitates a conceptual shift from viewing AZ scaffolds as static, homogeneous blocks to recognizing them as dynamic, tunable modules that are differentially deployed to shape the specific performance characteristics of diverse synaptic connections.

To understand why RIM is selectively essential at phasic MN-Is terminals, we must consider the distinct physiological strategies employed by these convergent inputs. MN-Ib and - Is are developmentally related glutamatergic motor neurons that express largely overlapping molecular machinery (Jetti et al., 2023; Nguyen et al., 2024), yet they achieve vastly different release set points (He et al., 2023; Newman et al., 2017, 2022). The mammalian AZ is a highly resilient, redundant structure where the loss of a single component rarely abolishes core release functions entirely (Acuna et al., 2016; Emperador-Melero & Kaeser, 2020; Kaeser et al., 2011; Südhof, 2012). Consistent with this resilience, we find that the total abundance of Ca_V_2 channels (CAC) is only modestly reduced in *rim* mutants at both inputs. Because tonic MN-Ib synapses rely predominantly on modulating Ca^2+^ influx to tune release (Chien et al., 2025; He et al., 2023; He & Dickman, 2025), they remain largely impervious to the loss of *rim*. In contrast, phasic MN-Is synapses depend heavily on a large, tightly coupled readily releasable vesicle pool to sustain high P_r_. Our nanoscale imaging reveals that RIM is uniquely enriched at MN-Is, where it positions the vesicle priming factor Unc13A in close proximity to Ca_V_2 channel nanodomains. This heavy reliance on RIM to maintain tight Ca_v_-vesicle coupling mirrors findings at the mammalian Calyx of Held, a giant, high-fidelity synapse where channel-vesicle topography governs release precision (Meinrenken et al., 2002) and RIM is essential to establish the tight coupling necessary for rapid, synchronous release (Han et al., 2011; Kaeser et al., 2011).

This input-specific reliance on RIM extends into homeostatic plasticity. Foundational studies previously identified RIM as an essential effector of presynaptic homeostatic potentiation (PHP), specifically driving the homeostatic expansion of the RRP (Müller et al., 2012). However, our ability to isolate inputs resolves a critical confound in these initial blended recordings: RIM is strictly required for the rapid expression of acute PHP at phasic synapses yet is entirely dispensable for chronic PHP at tonic synapses. Recent work has demonstrated that chronic PHP at MN-Ib is driven by enhanced Ca^2+^ influx, whereas acute PHP at MN-Is relies on expanding the RRP without altering overall Ca^2+^ influx (Chien et al., 2025). Thus, RIM’s selective requirement is consistent with the distinct mechanisms employed by these inputs.

Furthermore, we identify RIM as a key coordinator of homeostatic AZ compaction during acute PHP. Recent super-resolution studies have revealed that acute potentiation involves a striking structural condensation of Ca_V_2 channels and AZ scaffolds (Böhme et al., 2019; Ghelani et al., 2023; Mrestani et al., 2021, 2023). We find that this AZ compaction is entirely RIM-dependent. Recent biophysical studies have demonstrated that RIM and RIM-BP can undergo liquid-liquid phase separation (LLPS), condensing into a dense proteinaceous phase that can cluster Ca_V_2 channels and vesicle priming factors, providing a plausible mechanism for rapid AZ reorganization (Chen et al., 2020; Wu et al., 2019). By homeostatically condensing this nanostructure – perhaps by modulating the phase separation properties of a RIM condensate in response to acute signaling - and positioning Unc13A closer to Ca_V_2 channels, RIM could actively reorganize the lateral topography of the active zone, enabling recruitment of additional “loosely coupled” vesicles into the releasable pool, as supported by our EGTA sensitivity assays.

The specialization of RIM at the fly NMJ provides a useful physiological framework to interpret emerging evidence of AZ heterogeneity in the mammalian brain. While Drosophila uses a single *rim* gene to differentially tune convergent inputs, the mammalian genome encodes multiple *rim* genes and isoforms (e.g., RIM1α, RIM1β, RIM2) that exhibit synapse-specific expression and functions (Kaeser et al., 2008, 2012). This genetic expansion in mammals may have evolved in part to fulfill analogous tunable, input-specific roles to those we observe at the fly NMJ, potentially allowing mammalian circuits to independently set the P_r_, short-term plasticity, and long-term plasticity rules across billions of heterogeneous connections. For instance, while RIM is important for vesicle docking, priming, and Ca^2+^ channel tethering at classical fast synapses (Kaeser et al., 2011; Schoch et al., 2002), its structural organization and functional requirements differ substantially at dopaminergic terminals, which use a sparse, specialized active zone-like machinery to mediate neuromodulator release (Banerjee et al., 2022; C. Liu et al., 2018; Robinson et al., 2019). Furthermore, RIM1α is required for presynaptic long-term potentiation (LTP) at hippocampal mossy fiber synapses (Castillo et al., 2002), while its loss also affects the RRP (Schoch et al., 2002), underscoring a potentially conserved principle where synapses built for high-demand signaling or rapid plasticity rely more heavily on dedicated protein modules to meet those requirements. By deploying RIM proportionally to the functional demand for Ca_v_-vesicle coupling, diverse circuits may independently set their baseline P_r_ and dynamic range, perhaps in part by exploiting the biophysical tunability of LLPS condensates to adjust the local concentration and spatial organization of the release machinery across distinct synapse subtypes (Wu et al., 2025; Zhu et al., 2025).

Finally, our discovery that a core AZ scaffold like RIM acts as a tunable, input-specific module invites a re-evaluation of how we understand the active zone architecture more broadly. The AZ is composed of a conserved network of proteins, including ELKS/BRP, RIM-BP, Liprin-α, Piccolo/Bassoon/Fife, Syd-1, and Unc13 (Brockmann et al., 2020; Emperador-Melero & Kaeser, 2020; Haucke et al., 2011; Südhof, 2012). Much of the foundational literature characterizing mutants in these AZ genes at the Drosophila NMJ relies on blended recordings that may mask input-specific specializations similar to those we describe here. We anticipate that additional AZ proteins may exhibit input-specific, rather than strictly universal, functions when examined with comparable resolution. For instance, while RIM is specialized for the high-release phasic input, other components – such as specific Unc13 isoforms, RIM-BP, or BRP – may prove disproportionately important for the Ca^2+^ influx-dependent mechanisms of tonic MN-Ib synapses (Böhme et al., 2016; Ghelani et al., 2023; Pooryasin et al., 2021). Ultimately, active zones are not merely passive intermediaries situated between propagating action potentials and transmitter release. Rather, they are dynamic, computational hubs that actively participate in information transfer, using a modular architecture to diversify basal synaptic properties and sculpt complex plasticity across the billions of heterogeneous connections in the nervous system.

## MATERIALS AND METHODS

### Fly stocks

Drosophila stocks were raised at 25°C on standard molasses food. Unless otherwise specified, *w^1118^* served as the wild-type control and genetic background for all genotypes. For input-specific silencing, we crossed *UAS-BoNT-C* with Is-GAL4 (*GMR27E09-GAL4*) or Ib-GAL4 (*dHb9-GAL4*) (Han et al., 2022). We used the endogenously tagged *cac^sfGFP^* to visualize CAC (Gratz et al., 2019), and *GluRIIA^PV3^* mutants to induce chronic PHP (Han et al., 2023). All experiments utilized third-instar larvae of both sexes. See **Table S2** for a complete list of fly stocks, reagents, and software.

### Molecular biology

The *rim^ALFA^* allele was generated by WellGenetics (Taipei, Taiwan) via CRISPR/Cas9-mediated homology-directed repair (HDR). A guide RNA (gRNA) targeting the *rim* N-terminus near the start codon was cloned into a U6 promoter plasmid. A donor plasmid (pUC57-Kan backbone) was constructed to encode an ALFA tag (Götzke et al., 2019) inserted immediately after the ATG start codon, a 3xP3-DsRed selection cassette flanked by PiggyBac terminal repeats, and ∼1 kb upstream and downstream homology arms amplified from the injection strain. Silent PAM mutations were introduced into the donor to prevent Cas9 re-cleavage following HDR. The gRNA, hs-Cas9, and donor plasmids were co-injected into *w^1118^* embryos. F1 progeny were screened for 3xP3-DsRed fluorescence. Correct insertion at the *rim* locus was verified using junction PCR and Sanger sequencing. Donor backbone integration was excluded via integration-specific PCR. The PiggyBac-flanked selection cassette was subsequently excised, leaving a TTAA footprint between the ALFA tag and the *rim* coding sequence; excision was confirmed by PCR and sequencing.

### Electrophysiology

All dissections and sharp-electrode current clamp recordings were performed as previously described (Han et al., 2023). Preparations were bathed in a modified hemolymph-like saline (HL-3) containing: 70mM NaCl, 5mM KCl, 10mM MgCl_2_, 10mM NaHCO_3_, 115mM Sucrose, 5mM Trehelose, 5mM HEPES, and 0.4mM CaCl_2_ (pH 7.2). The internal organs, brain, and ventral nerve cord were removed prior to recording. Recordings were acquired on an Olympus BX61 WI microscope equipped with a 40x/0.80 NA water-dipping objective, using an Axoclamp 900A amplifier, Digidata 1440A acquisition system and pClamp 10.5 software (Molecular Devices). Data were analyzed only from cells with initial resting potentials more hyperpolarized than −60 mV and input resistances >5 MΩ. All recordings were performed on abdominal muscle 6 or 7 in segments A2, A3, or A4 of third-instar larvae.

Miniature excitatory postsynaptic potentials (mEPSPs) were recorded for 60 secs in the absence of any stimulation and analyzed using MiniAnalysis (Synaptosoft) and Excel (Microsoft) software. The average mEPSP amplitude for each NMJ was calculated from approximately 100 consecutive events. Excitatory postsynaptic potentials (EPSPs) were evoked by delivering 20 stimuli (0.5 msec duration, 0.5 Hz) to the motor innervation via an ISO-Flex stimulus isolator (A.M.P.I.) at intensities required to induce single EPSPs.

Two-electrode voltage-clamp (TEVC) recordings were performed as previously described (Chien et al., 2025). Larvae were bathed in HL-3 containing the indicated CaCl_2_ concentrations. Recordings were acquired using an Olympus BX61WI microscope (40x/0.8 NA water-dipping objective) and an Axoclamp 900A amplifier (Molecular Devices) (He et al., 2023). We recorded from abdominal muscle 6 (segments A3 and A4), requiring an initial resting potential below −60 mV and an input resistance >5 MΩ. mEPSPs were recorded for 60 secs.

The average mEPSP amplitude was calculated from approximately 100 events per recording. Excitatory postsynaptic currents (EPSCs) were evoked by delivering 20 stimuli (0.5 msec duration, 0.5 Hz) via an ISO-Flex stimulus isolator. To acutely block postsynaptic receptors and induce PHP, semi-dissected larvae (dorsal incision only) were incubated in HL-3 containing 20 µM philanthotoxin-433 (PhTx; Sigma) for 12 min (Kiragasi et al., 2017). Preparations were then washed three times with HL-3 prior to recording. To estimate the readily releasable pool (RRP) size, we evoked EPSCs with a 30-stimulus train (60 Hz) in 3 mM Ca^2+^ HL-3. A linear fit was applied to the cumulative EPSC data (stimuli 19–30) and back-extrapolated to time 0; this Y-intercept value was divided by the mean mEPSP amplitude to estimate RRP size. To evaluate EGTA sensitivity, we incubated larval fillets in 0 Ca^2+^ HL-3 supplemented with 50 μM EGTA-AM (Sigma-Aldrich) for 10 mins, followed by three washes with standard HL-3 prior to recording (Chien et al., 2025). Where indicated, EGTA-AM was applied following 10 min PhTx incubation.

### Immunocytochemistry

Third-instar larvae were dissected in ice-cold 0 Ca^2+^ HL-3 and immunostained as previously described (Kikuma et al., 2017). Preparations were fixed in ice-cold 100% methanol (5 min), Bouin’s fixative (5 mins), or 4% PFA (10 mins), followed by three 10 min washes in PBST (PBS + 0.1% Triton X-100). Samples were blocked in 5% Normal Donkey Serum, incubated with primary antibodies overnight at 4°C, washed 3x in PBST, and incubated with secondary antibodies for 2 hours at room temperature. Following three final PBST washes, samples were equilibrated in 70% glycerol and mounted in VectaShield for confocal imaging or ProLong Glass (ThermoFisher) for STED microscopy. Native Cac^GFP^ was imaged without secondary amplification in confocal experiments. Primary antibodies included: mouse anti-BRP (nc82, 1:200), mouse anti-GFP (1:100), guinea pig anti-RBP (1:2000), rabbit anti-Unc13A (1:1000), guinea pig anti-Unc13B (1:500), rabbit anti-ALFA (1:500), and Alexa Fluor 647-conjugated goat anti-HRP (1:400). Secondary antibodies included STAR RED-conjugated secondaries for STED imaging (1:200) and Cy3-, AF488-, AF594-, or AF647-conjugated secondaries (1:400). See Table S2 for complete details and sources. To ensure robust PhTx-induced remodeling for imaging experiments, we utilized a modified protocol employing 80 µM PhTx for 15 mins. Preparations were co-stained with BRP to verify consistent remodeling.

### Confocal imaging and analysis

Confocal images were acquired on a Nikon A1R microscope (100x APO 1.40NA oil objective) using NIS Elements software and 405, 488, 561, and 647 nm laser lines (Kiragasi et al., 2020). For intensity quantification of BRP, RBP, CAC, Unc13A, Unc13A, and RIM, we acquired z-stacks (0.150 µm z-step, 40 nm x/y pixel) using identical gain and laser settings across all samples within an individual experiment. Nikon Elements software was utilized to identify individual puncta within maximum intensity projections of boutons and to calculate mean fluorescence intensities. The average fluorescence intensity was expressed as a single number per bouton. All confocal measurements derived from M6/7 terminal boutons (1 bouton per Ib NMJ; 1-3 boutons per Is NMJ) across >10 NMJs from >4 animals.

### STED imaging and analysis

Stimulated Emission Depletion (STED) microscopy was performed as previously described using an Abberior STEDYCON system mounted on a Nikon Eclipse FN1 microscope (He et al., 2023). The system used 640, 561, 488, and 405 nm excitation lasers, a 775 nm pulsed STED laser, and three avalanche photodiode detectors. Multichannel 2D STED images (1-2 boutons per field of view) were acquired with a 100x Nikon Plan APO 1.45 NA oil objective (15 nm pixel size, 10 µs dwell time, 15x line accumulation). Abberior STAR Red and Alexa Fluor 594 dyes were depleted at 775 nm. Time-gating was set to 1 nsec (6 nsec width), and photons were detected sequentially (STAR RED: 675 nm; Alexa Fluor 594: 600nm). Raw STED images were deconvolved using SVI Huygens software (default settings, theoretical STED PSF). Protein-covered areas were quantified from raw images using the NIS Elements general analysis toolkit. We manually selected active zones displaying optimal planar orientation and determined protein area and equivalent diameter by applying a fixed intensity threshold to the 600 nm and 675 nm channels. BRP and RIM nanomodules were quantified using local maxima detection in ImageJ. All STED measurements exclusively evaluated 2D images of M6 NMJs (segments A3/4) from >4 animals.

Deconvolved 16-bit STED images were used to quantify distances between protein puncta via line-profile estimation of peak-to-peak distances (Pooryasin et al., 2021). Only well-defined puncta located within the same focal plane were included in the analysis. Planar or side-view synapses displaying clear protein signals within the same z-plane were manually traced using the ImageJ line profile tool (1 pixel thickness). Fluorescence intensity profiles were extracted for each channel, and the positions of maximal intensity for each protein were determined. Inter-protein distances were calculated as the peak-to-peak distance in pixels and subsequently converted to nanometers.

### Statistical analysis

Data were analyzed using GraphPad Prism (v8.0), MiniAnalysis, SVI Huygens Essential (v22.10), and Microsoft Excel (v16.22). Normality was assessed using the D’Agostino & Pearson omnibus test. Normally distributed data were compared via an unpaired or paired 2-tailed Student’s t-test (with Welch’s correction) or a one-way ANOVA incorporating Tukey’s multiple comparison test. Non-normally distributed data were evaluated using the Mann-Whitney test. In all figures, error bars indicate the standard error of the mean (SEM), and significance is denoted as: \**p*<0.05, \*\**p*<0.01, \*\*\**p*<0.001, \*\*\*\**p*<0.0001; ns=not significant. Detailed statistical information for all datasets is provided in **Table S1**.

## AUTHOR CONTRIBUTIONS

R.S. and D.D. designed the research. R.S. generated reagents, performed most experiments, and analyzed the data. P.P. and C.D. performed genetic experiments and W.D. validated reagents. The manuscript was written by R.S. and D.D.

## ACKNOWLEDGEMENTS

We acknowledge the Developmental Studies Hybridoma Bank (Iowa, USA) for antibodies and the Bloomington Drosophila Stock Center for fly stocks (NIH P40OD018537). This work was supported by an NSF GRSF fellowship to R.S. (DGE-1842487) and a grant from the National Institutes of Health to D.D. (NS126654).

---

**Supplemental Figure S1:**
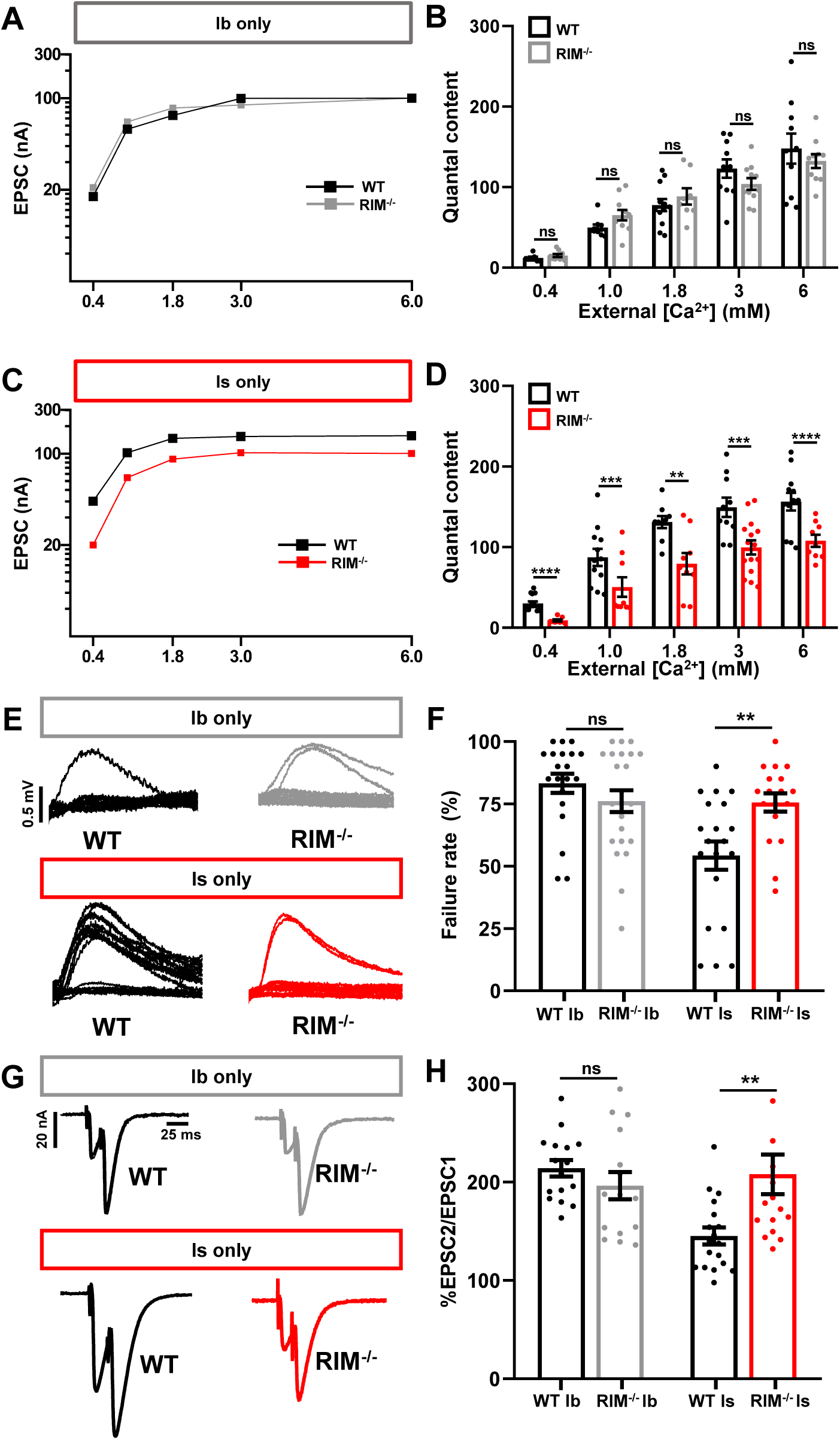
Additional electrophysiological analyses confirm loss of *rim* only impacts transmission at MN-Is. **(A,B)** Plot of EPSC values as a function of external [Ca^2+^] (A) and quantification of quantal content (B) for isolated MN-Ib in wild-type and *rim*^-/-^synapses. No differences are observed in *rim* mutants. **(C,D)** Corresponding data for isolated MN-Is, revealing clear reductions in *rim* mutants. **(E,F)** Representative traces (E) and quantification of failure rate (F) across 20 stimuli (0.5 Hz) at 0.10 mM extracellular [Ca^2+^], confirming lowered release probability selectively at *rim*^-/-^ MN-Is. **(G,H)** Representative paired-pulse EPSC traces (G) and quantification of paired-pulse ratios (EPSC2/EPSC1) (H) at 1.8 mM extracellular [Ca^2+^] (16.7 msec interstimulus interval), revealing deficits in release probability only at *rim^-/-^* MN-Is. Error bars indicate ±SEM. Additional statistical details are provided in Table S1.

**Supplemental Figure S2:**
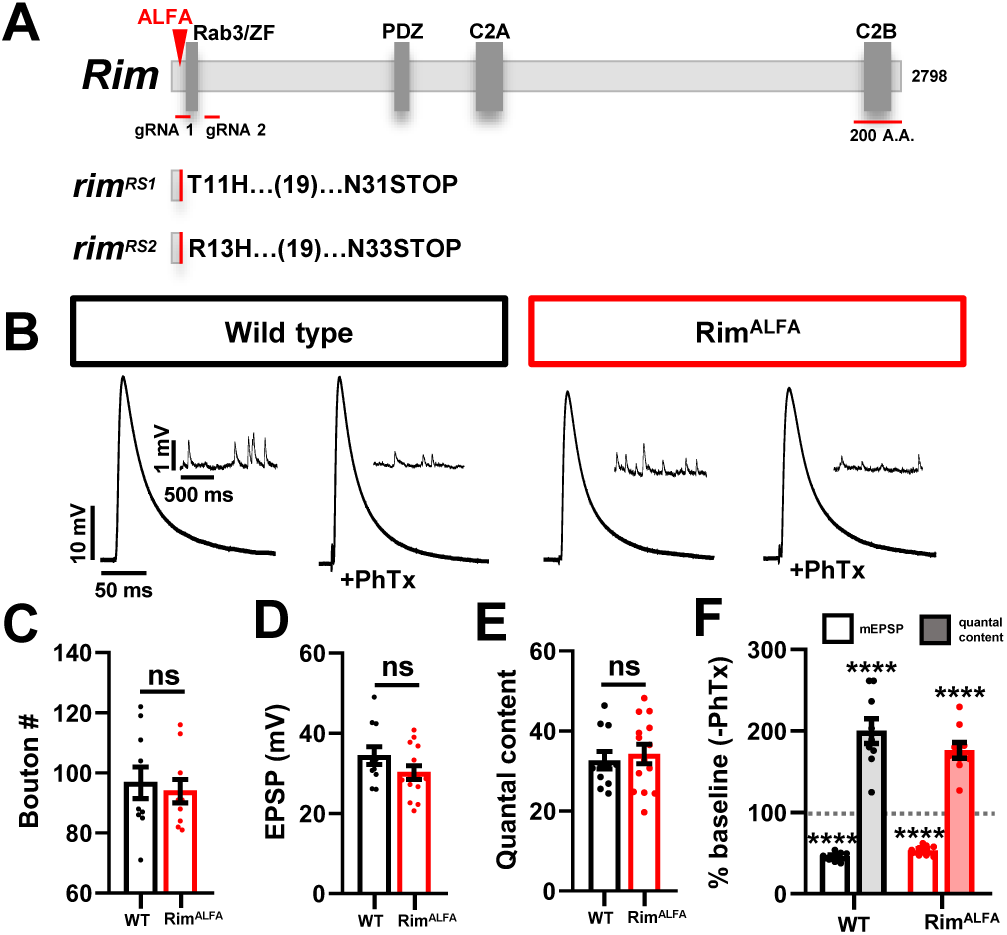
Synaptic transmission, growth, and plasticity are not perturbed in *rim^ALFA^*. **(A)** Schematic of the Drosophila *rim* locus indicating sgRNA targets for CRISPR mutagenesis and the ALFA tag insertion site, along with the resulting mutant alleles (*rim^RS1^ and rim^RS2^*) and early stop codons. **(B)** Representative mEPSP and EPSP traces at baseline and following PhTx application. **(C-F)** Quantification of bouton number (C), EPSP amplitude (D), and quantal content values (E,F), revealing no changes in synaptic growth, transmission, or presynaptic homeostatic potentiation in *rim^ALFA^* animals. Error bars indicate ±SEM. Additional statistical details are provided in Table S1.

